# Connexin-43 dependent ATP release mediates macrophage activation during peritonitis

**DOI:** 10.1101/424333

**Authors:** Michel Dosch, Joël Zindel, Fadi Jebbawi, Nicolas Melin, Daniel Sanchez-Taltavull, Deborah Stroka, Daniel Candinas, Guido Beldi

## Abstract

Peritonitis is the consequence of bacterial spillage into a sterile environment by gastrointestinal hollow-organ perforation that may lead to fulminant sepsis. Outcome of peritonitis-induced sepsis critically depends on macrophage activation by extracellular ATP release and associated para- and autocrine signaling via purinergic receptors. Mechanisms that mediate and control ATP release, however, are poorly understood. Here we show that TLR-2 and -4 agonists trigger ATP release via Connexin-43 (CX43) hemichannels in peritoneal macrophages leading to poor survival during sepsis. In humans, CX43 expression was upregulated on macrophages isolated from the peritoneal cavity in patients with intraperitoneal infection but not in healthy controls. Using a murine caecal ligation and puncture (CLP) model, we identified increased CX43 expression in activated infiltrating peritoneal, hepatic and pulmonary macrophages. Conditional MAC-CX43 KO *Lyz2*^cre/cre^*Cx43*^flox/flox^ mice were developed to specifically assess the CX43 impact in macrophages. Both macrophage-specific CX43 deletion (using *Lyz2*^cre/cre^*Cx43*^flox/flox^ mice) or pharmacological CX43 blockade were associated with reduced cytokine secretion by macrophages in response to LPS and CLP. This was ultimately resulting in increased survival in *Lyz2*^cre/cre^*Cx43*^flox/flox^ mice and after pharmacological blockade. Specific inhibition of the purinergic receptor P2RY1 abrogated CX43 elicited cytokine responses. In conclusion, inhibition of autocrine ATP signaling via CX43 on macrophages and P2RY1 improves sepsis outcome in experimental peritonitis.

**Brief Summary:** Connexin-43-mediated ATP release from macrophages in response to TLR-4 and -2 agonists modulates autocrine activation of macrophages in a P2Y1-dependent manner, ultimately determining sepsis survival.

## INTRODUCTION

Sepsis is associated with high mortality and was now recognized as a Global Health Priority by the World Health Organization (1). Mechanistically sepsis is characterized by a dysregulation of the host immune response to bacteria resulting in local and systemic exacerbated pro-inflammatory response and a concomitant protracted anti-inflammatory response (2, 3). Accordingly, immuno-regulatory drugs have been considered as a potential treatment, however, clinical results have been unsatisfactory so far (4). Despite the urgency to improve patient outcomes and major advances in the understanding of sepsis, a specific therapy is still lacking.

Peritoneal sepsis, as a result of a hollow-organ perforation, is typically associated with invasion of the peritoneal cavity by intestinal contents containing gastrointestinal bacteria. Gastrointestinal commensals and pathogens proliferate in the peritoneal cavity and release pathogen associated molecular patterns that interact with pathogen recognition receptors on local macrophages and neutrophils to initiate strong inflammatory reactions (5).

Macrophage activation is critically modulated by extracellular ATP and associated purinergic signaling via autocrine/paracrine mechanisms (6). During sepsis, purinergic signaling regulates core inflammatory cell functions such as migration and inflammatory mediators production (6–8). ATP depletion or blockade of purinergic receptors were shown to increase survival and improve outcome of sepsis in rodent models (9, 10). The initiating event triggering purinergic signaling is the release of extracellular nucleotides and includes active vesicular and channel-mediated release in addition to the passive release from necrotic cells (11). Active ATP release from inflammatory cells can occur both via vesicular exocytosis or via connexin or pannexin hemichannels, mainly connexin-43 (CX43) and pannexin-1 (11, 12). In sepsis, it is still unknown which mechanisms mediate ATP release and if targeting these mechanisms would impact outcomes. In addition, it remains to be described which inflammatory cells are involved in ATP release and to what extend ATP determines autocrine/paracrine inflammatory cell regulation during sepsis.

By screening release mechanisms of ATP using small molecule blockers, we have identified CX43 to be critical in peritoneal sepsis. CX43 is a gap junction protein forming either gap junctions or hemichannels and has been shown to mediate ATP release from inflammatory cells including macrophages (10, 13). CX43 full knockout is perinatally lethal in mice due to cardiac malformation (14, 15). Thus, in order to specifically explore the function of CX43 on inflammatory cells, we developed a conditional MAC-CX43 KO mouse (*Lyz2*^cre/cre^*Cx43*^flox/flox^), in which CX43 is specifically deleted on macrophages and neutrophils. We identified that peritoneal macrophages express high CX43 levels in mice and humans and that CX43 expression on macrophages critically controls ATP release, local and systemic inflammation, and ultimately survival in a model of abdominal peritonitis via the purinergic receptor P2Y1. Our findings suggest that CX43 represents a new potential target for sepsis therapy.

## RESULTS

*Small peritoneal macrophages release extracellular ATP via Connexin-43 during peritonitis*. Inflammatory cell populations present in the peritoneal cavity during peritonitis were characterized using CLP as a murine model of peritoneal sepsis and include infiltrating small peritoneal macrophages (SPM) and resident large peritoneal macrophages (LPM) (16, 17). Ten hours after CLP, SPM populated the peritoneal cavity while the fraction of LPM disappeared (**Figure 1 A-B, Supplemental Figure 1 A-B**). Additionally, two large populations of neutrophils invaded the peritoneal cavity upon sepsis while dendritic cells disappeared. Given the observation that SPM specifically populate the peritoneal cavity after CLP, we tested the ability of these macrophages to secrete ATP. Isolated C57BL/6 wild-type (WT) SPM actively released ATP in response to TLR-4 agonist LPS (1μg/ml) as well as the TLR-2 agonist Pam3CSK4 in a dose-dependent manner (**Figure 1 C-D**). ATP released in the extracellular space was promptly degraded by ecto-nucleotidases, as lower levels of extracellular ATP were observed in absence of treatment with ARL67156 trisodium salt, an ecto-ATPase inhibitor (**Supplemental Figure 1 C**).

**Figure 1.**
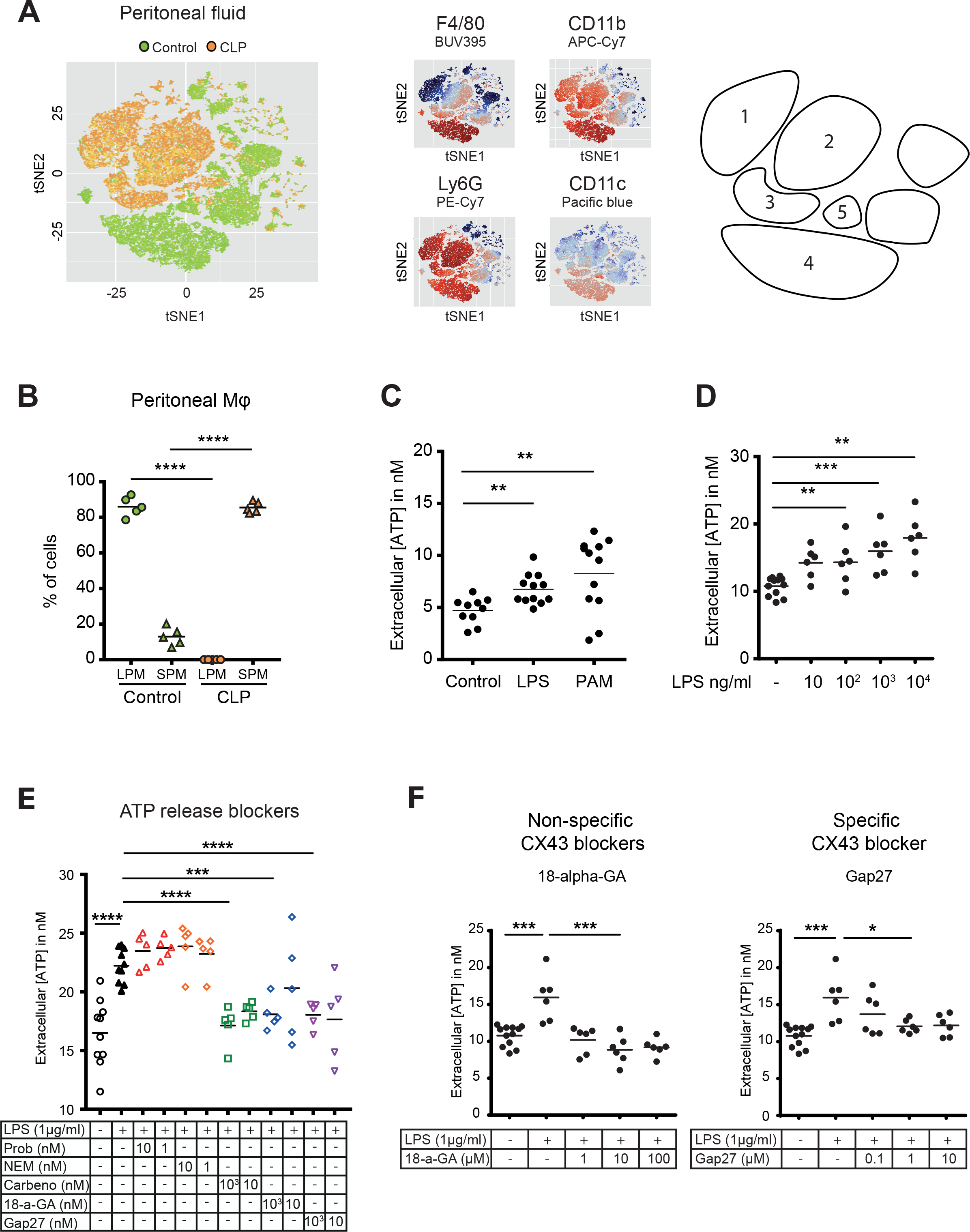
Small peritoneal macrophages release ATP during sepsis via Connexin-43 hemichannels. (**A**) Intraperitoneal cell fractions characterized by flow cytometry in controls (green) and 10 hours after caecal ligation and puncture (CLP, orange) in a tSNE plot of viable (AmCyan^low^) CD45^high^ cells. Populations were defined based on F4/80 (BUV395), CD11b (APC/Cy7), Ly6G (PE-Cy7) and CD11c (Pacific blue). In response to CLP, elevated levels of neutrophils (**1**)/(**2**) (F4/80^low^, CD11b^high^, Ly6G^high^) and small (infiltrating) peritoneal macrophages (**3**) (F4/80^int^, CD11b^int^) were observed but not large (resident) peritoneal macrophages (**4**) (F4/80^high^, CD11b^high^) or dendritic cells (**5**) (F4/80^low^, CD11b^low^, Ly6G^low^, CD11c^high^). (**B**) Relative frequency of large peritoneal macrophages (LPM) and small peritoneal macrophages (SPM) in the peritoneal cavity of controls (green) and 10 hours after CLP (orange) (unpaired t-test). (**C**) Extracellular ATP levels in the supernatant of primary peritoneal macrophages after 30 minutes stimulation with LPS (TLR-4 agonist) or Pam3CSK4 (PAM, TLR-2 agonist). Cells were isolated from C57 Bl/6 WT mice (unpaired t-test). (**D**) Dose-dependent ATP release by peritoneal macrophages in response to LPS as quantified by a luciferin-luciferase assay (unpaired t-test). (**E**) LPS-induced ATP release by macrophages can be blocked by carbenoxolone ((Carbeno) combined pannexin/connexin blockade), 18-alpha-GA (18-a-GA) global connexin blockade), Gap27 (specific blockade of connexin 43 (CX43)) but not by probenecid ((Prob) pannexin channel blocker) and N-Ethylmaleimide ((NEM) blockade of vesicular exocytosis) (unpaired t-test). (**F**) Non-specific (18-a-GA) and specific (Gap27) blockade of connexin hemichannels decreases LPS-induced ATP release from primary murine WT peritoneal macrophages in a dose-dependent manner (unpaired t-test).

To screen cellular mechanisms underlying LPS-induced ATP release, specific blockers of (hemi)channels and vesicular exocytosis were administrated (**Figure 1 E**). LPS-induced ATP release was blocked by carbenoxolone (combined pannexin/connexin blockade), 18-alpha-GA (global connexin blockade) and Gap27 (blockade of Connexin-43 (CX43)) but not by probenecid (pannexin channel blockade) and N-Ethylmaleimide (blockade of vesicular exocytosis) revealing a specific role of CX43 (**Figure 1 E**). The relevance of CX43 is substantiated by dose-dependent unspecific (18-alpha-glycyrrhetinic acid) and specific (Gap27) CX43 blockade (**Figure 1 F**).

*Connexin-43 expression is induced on macrophages in a MyD88/TRIF dependent manner*. In humans, fluids obtained from peritoneal lavage during abdominal operations were analysed by flow cytometry. CX43^high^ macrophages were observed in the peritoneal fluid of patients with peritonitis but were not detected in patients without signs of intra-abdominal infection (**Figure 2 A, Supplemental Figure 2 A-C, Supplemental Table 1**).

**Figure 2.**
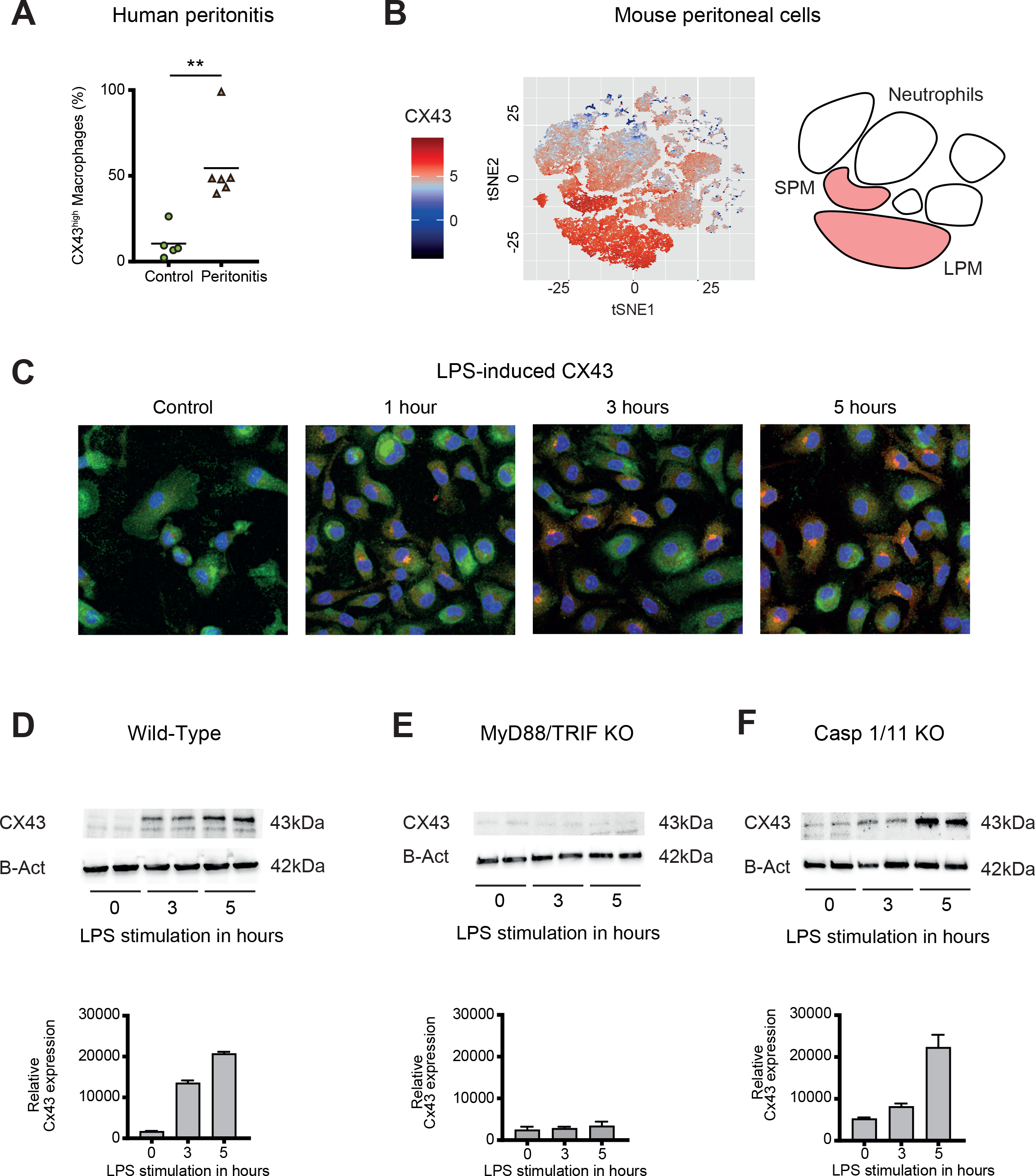
Connexin-43 expression is induced on macrophages in a MyD88/TRIF dependent manner. (**A**) Percentage of CX43^high^ macrophages in the peritoneal fluid collected from control patients without peritoneal inflammation (green) compared to patients with peritonitis (orange). (**B**) Specific CX43 expression (Alexa Fluor 488) in macrophages (small and large peritoneal macrophages, SPM and LPM) among peritoneal viable (AmCyan^low^) CD45^high^ cells. (**C-D**) Increased expression of CX43 in macrophages following stimulation with LPS (1 µg/ml) for the indicated time-points (PE-Texas red: CX43; FITC: F4/80; blue: DAPI). (**E-F**) Abrogated CX43 protein expression in primary peritoneal macrophages from MyD88/TRIF double KO mice but not from caspase 1/11 double KO mice upon stimulation with LPS 1 μg/ml. Quantification of immunoblots was made using ImageJ, unpaired t-test.

In mouse, analysis of cellular fractions present in the peritoneal cavity during peritonitis revealed that CX43-dependent ATP release is macrophage-specific, since both SPM and LPM are CX43-positive, while neutrophils are CX43-negative (**Figure 2 B**). CX43 protein expression in primary WT murine macrophages is induced by LPS (**Figure 2 C-D**). Mechanistically, MyD88 and TRIF are required for the expression of CX43 after LPS stimulation (**Figure 2 E**), while it is independent of caspase-1 and caspase-11 (**Figure 2 F**). Thus, CX43 in SPM is responsible for LPS-induced ATP release in peritoneal sepsis in a MyD88/TRIF dependent manner.

*Connexin-43 expressing macrophages are recruited systemically during peritonitis*. Sepsis is a systemic inflammatory reaction that leads to extensive organ damage remote from the primary infection site, typically in the liver and the lungs. To understand whether CX43 positive macrophages would be present locally at the site of infection only or in remote organs as well, we analysed hepatic and pulmonary tissues in our CLP model. In the liver of controls, CX43 was expressed on cholangiocytes only, while after CLP it was highly upregulated on non-parenchymal cells in a time-dependent manner (**Figure 3 A-B**). CX43 protein expression in the liver was dependent on MyD88/TRIF activation (**Figure 3 C**). Following CLP, infiltrating monocytes/macrophages and neutrophils increased in the liver while resident macrophages (Kupffer cells) remained constant (**Figure 3 D-E**). CX43 expression in the liver was restricted to infiltrating monocytes/macrophages, while neutrophils and resident macrophages were CX43 low (**Figure 3 F**). Hepatic lympho-cellular populations, such as T cells, B cells, and NK cells did not express CX43 (**Supplemental Figure 2 D-F).**

**Figure 3.**
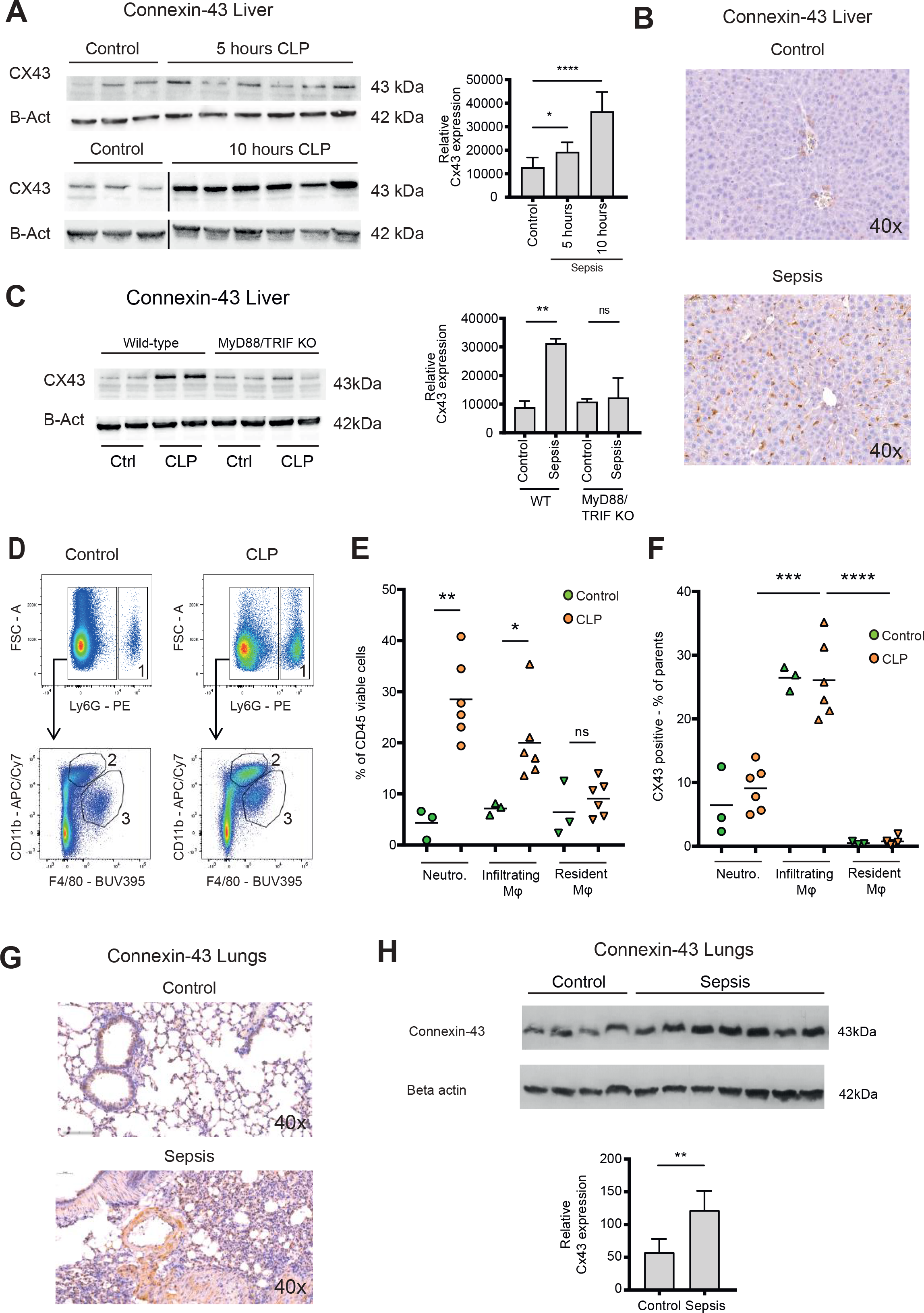
Connexin-43 expressing macrophages are recruited systemically during sepsis. (**A**) Elevated CX43 protein expression in the liver 5 and 10 hours after CLP compared to controls. (**B**) Elevated expression of CX43 on non-parenchymal cells in the liver 10 hours after CLP compared to controls. (**C**) Abrogated CX43 protein expression in the liver of MyD88/TRIF double KO mice 10 hours after CLP. (**D-E**) Following CLP (orange), infiltrating monocytes/macrophages (**2**) and neutrophils (**1**) increased in the liver, while resident macrophages (**3**) remained constant compared to controls (green), unpaired t-test. (**F**) Infiltrating monocytes/macrophages expressed high CX43 levels while neutrophils (**1**) and resident macrophages (**3**) were CX43 low. (**G, H**) Elevated expression of CX43 on non-parenchymal cells in the lungs in response to CLP compared to control mice, unpaired t-test.

In the lungs, CX43 is constitutively expressed on muscle cells and resident macrophages populations (**Figure 3 F**). In the acute respiratory distress syndrome that is associated with severe peritonitis, alveolar infiltration of CX43 inflammatory cells was observed (**Figure 3 G, H).** These results reveal that CX43 is specifically expressed in the liver and lungs as remote target organs during sepsis.

*Connexin-43-mediated local and systemic ATP release contributes to macrophage over-activation via P2RY1*. Levels of extracellular ATP were increased in the peritoneal cavity as well as in the systemic circulation 10 hours after CLP (**Figure 4 A-B**). To understand the importance of CX43 in ATP release from macrophages and its impact on macrophage activation, we developed conditional MAC-CX43 KO (*Lyz2^cre/cre^, Cx43^flox/flox^*) mice in which CX43 is specifically deleted in activated macrophages (**Supplemental Figure 3 A-E**). Following CLP, we observed lower systemic ATP levels in MAC-CX43 KO mice compared to the WT mice, suggesting a role for CX43 in systemic ATP release, supporting the results in figure 1 showing that local ATP release from macrophages is CX43-dependent (**Figure 4 B**). In a next step, we assessed autocrine responses of CX43-dependent ATP release in primary peritoneal macrophages. Pharmacological blocking (Gap27) or genetic deletion of CX43 decreased pro-inflammatory cytokines levels (TNF-alpha and IL-6) 3 and 6 hours post-LPS stimulation in primary peritoneal macrophages in vitro (**Figure 4 C-D**). Next, we aimed to understand whether CX43 blocking or deletion, and consequent decrease in ATP release, regulate macrophage differentiation. Functional markers of M1 differentiation, such as *Inos* and *Il12rb*, were decreased in macrophages under CX43 blocking or deletion (**Supplemental Figure 4 A-B**), while markers of M2 differentiation, including *Arg1, Tgfb* and *Il-10*, were comparable among groups (**Supplemental Figure 4 C-E**). Phagocytosis of IgG coated latex beads was not impacted by CX43 blocking or deletion (**Supplemental Figure 3 F**). Therefore, autocrine CX43-dependent ATP signalling alters cytokine secretion and M1 macrophage differentiation but not phagocytosis.

**Figure 4.**
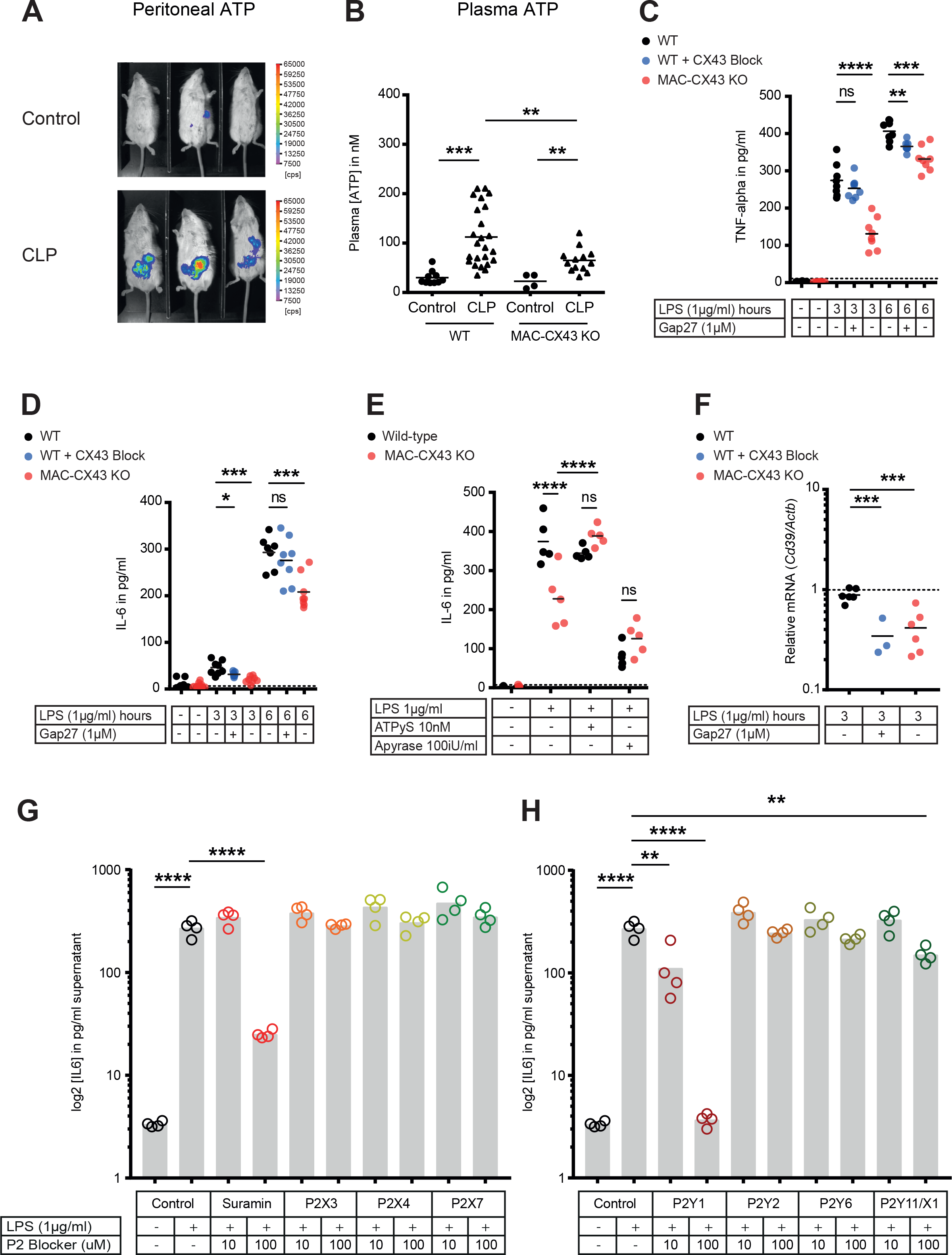
Connexin-43-mediated local and systemic ATP release contributes to macrophage over-activation via P2RY1. (**A**) Extracellular ATP levels as assessed by luminescence of i.p. injected 5×10^6^ HEK293-pmeLUC D-luciferin activated cells (in counts per second (cps)). Luminescence was significantly higher 10 hours after CLP compared to sham-operated controls. (**B**) Direct measurement of extracellular ATP in the plasma collected from vena cava inferior is elevated in response to CLP and higher in WT mice compared to MAC-CX43 KO mice. Each dot is representative for a single animal (unpaired t-test). (**C-D**) Inhibition (Gap27 (1 µM)) or genetic deletion of CX43 decreased TNF-alpha (**C**) und IL-6 (**D**) production from primary peritoneal macrophages in response to stimulation with LPS (1 μg/ml) for 3 and 6 hours. (**E**) IL-6 release was restored by activation of purinergic receptors by exogenous administration of ATPуS (10 nM) and abrogated in response to apyrase (100 IU/ml) (unpaired t-test). (**F**) *Cd39* mRNA levels decreased in response to CX43 blockade or deletion. (**G-H**) IL-6 release from primary peritoneal macrophages in response to LPS (1 µg/ml) and by blocking major P2X (**G**) and P2Y (**H**) purinergic receptors.

The administration of ATPgammaS, a non-hydrolyzable form of ATP, reverted the downstream effects of CX43 blockade and abrogated ATP release by restoring IL-6 and TNF-alpha levels in macrophages after CX43 blockade or deletion to control levels (**Figure 4 E, Supplemental Figure 4 G**). Administration of apyrase, a soluble ecto-ATPase consuming extracellular ATP, decreased pro-inflammatory cytokines levels. Thus, the downstream effects of LPS-induced CX43 -dependent ATP release is mediated via purinergic receptors. To identify the specific P2 receptor responsible for this effect, a screen using different P2 receptors blockers was performed to explore the effects of LPS-dependent TNF-alpha and IL-6 release from macrophages (**Supplemental Table 2**). The observed reversal of TNF-alpha and IL-6 secretion following unspecific P2 receptor blockade (suramin) and specific P2RY1 blockade (MRS 2279) indicate a crucial role of P2RY1 in macrophages activation following LPS stimulation (**Figure 4 G-H, Supplemental Figure 5 A-B**). Gene expression of purinergic ATP receptors (P2X and P2Y receptors) were not differently regulated between MAC-CX43 KO and WT (**Supplemental Figure 5 C**). Extracellular ATP is hydrolysed by the ecto-nucleotidase CD39 to ADP/AMP and by ecto-5’nucleotidase CD73 to adenosine. In response to MAC-CX43 KO or blockade, CD39 mRNA levels were reduced in MAC-CX43 KO macrophages compared to WT controls, while no difference between WT and MAC-CX43 KO was observed for ecto-5’nucleotidase CD73 (**Figure 4 F, Supplemental Figure 4 H**). Taken together, CX43 blockade or deletion reduce ATP secretion and its autocrine downstream effects on macrophages via P2RY1.

*Improved survival and decreased local and systemic cytokine production in response to connexin-43 blocking or deletion during abdominal sepsis*. To test CX43 impact on sepsis outcome, we used our MAC-CX43 KO mice (*Lyz2^cre/cre^, Cx43^flox/flox^*) and compared them to mice heterozygous for lyzozyme 2 cre sequence as controls (*Lyz2^cre/wt^, Cx43^flox/flox^*) (**Supplemental Figure 3 A-E**), pharmacological inhibition with Gap27 and controls. Murine sepsis score (**Supplemental Table 3**), weight loss, survival time, bacterial load, and markers of inflammation were monitored. The extent of sepsis, as assessed by the murine sepsis score, was significantly reduced in MAC-CX43 KO compared to WT mice without a difference in weight loss (**Figure 5 A-B**). CLP is a lethal model with a survival time expected to be reached between 18 and 24 hours following surgery. CX43 blocking or specific genetic deletion in CLP operated mice was associated with significantly prolonged survival time compared to control WT C57BL/6 mice (**Figure 5 C**). Bacterial load was not different in both peritoneal fluid and blood excluding altered phagocytosis by macrophages to be relevant (**Supplemental Figure 6 A-B**).

**Figure 5.**
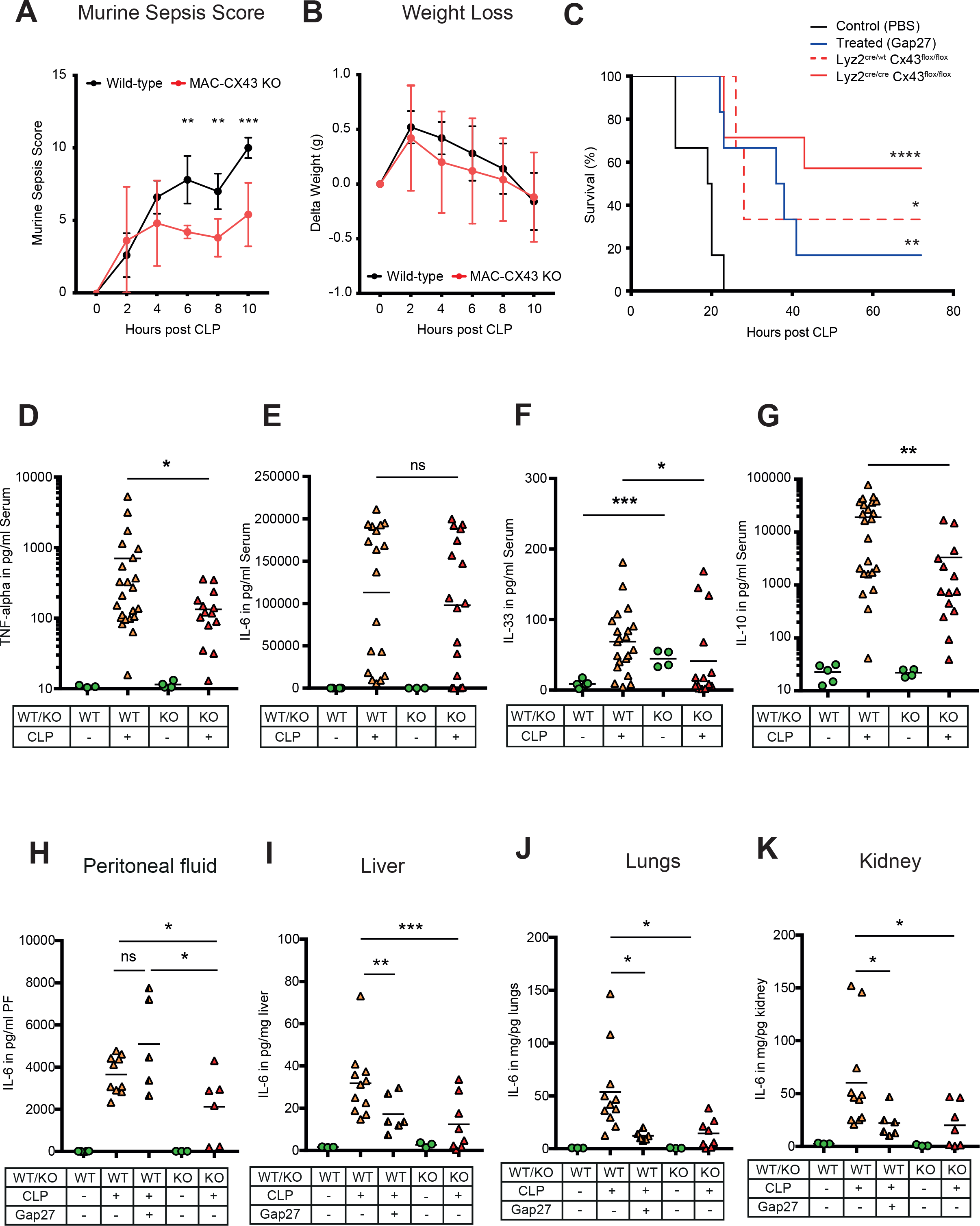
Improved survival and decreased local and systemic cytokine production in response to connexin-43 blocking or deletion during abdominal sepsis. (**A-B**) Clinical outcome using Murine Sepsis Score and weight loss following CLP. (**C**) Survival of mice following caecal ligation and puncture (CLP). Mice treated with Gap27 (blue), a specific CX43 blocker, homozygous (*Lyz2^cre/cre^*, *Cx43^flox/flox^*, red) and heterozygous cre (*Lyz2^cre/wt^*, *Cx43^flox/flox^*, red dashed) were compared to non-treated WT controls (black) (Log-rank (Mantel-Cox) test). (**D-G**) Systemic levels of TNF-alpha, IL-6, IL-33 and IL-10 in the serum of WT and MAC-CX43 KO mice operated with CLP. (**H-K**) IL-6 in peritoneal fluid (PF) serum, lungs, kidney and liver. Levels are expressed in pg/mg tissue respectively pg/ml of serum or peritoneal fluid.

In order to assess if cellular fractions change in MAC-CX43 KO mice compared to WT mice during sepsis, peritoneal cells were characterized by mass and flow cytometry. At baseline, two resident macrophages clusters were identified (**Supplemental Figure 7 A-B**). The number of CX43^high^ resident macrophages was higher in the peritoneal cavity of WT mice compared to MAC-CX43 KO mice at baseline, which might be due to low Lyz2 promoter activation during their maturation (**Supplemental Figure 7 B**). Resident macrophages disappeared 10 hours after CLP and neutrophils and different clusters of infiltrating monocytes/monocyte-derived macrophages clusters were increased in the peritoneal cavity (**Supplemental Figure 7 C**). However, no difference in cell numbers was observed between WT and MAC-CX43 KO mice, indicating that even though macrophages were differently activated, their numbers were not impacted by CX43 deletion or blockade. Relative fractions of CD4 and CD8 T cells decreased in septic animals compared to control without difference between WT and MAC-CX43 KO (**Supplemental Figure 8 A-B**).

Inflammatory cytokines were assessed systemically (serum), locally (peritoneal fluid), and in key organs typically injured during sepsis (lungs, liver, kidney, intestine). We observed decreased serum levels of pro-inflammatory cytokines (TNF-alpha, IL-6, IL-33) in MAC-CX43 KO compared to WT mice 10 hours after CLP (**Figure 5 D-F**). In addition, IL-10 levels were lower, indicating a decrease of both pro- and anti-inflammatory cytokines in MAC-CX43 KO mice (**Figure 5 G**). In the peritoneal fluid and in the peripheral effector organs, we observed decreased IL-6 levels (with the exception of the intestine) of mice under CX43 blocking or deletion compared to non-treated WT mice (**Figure 5 H-K, Supplemental Figure 6 C**). A relevant role of neutrophils can be excluded, first because of limited expression of CX43 in response to sepsis (**Figure 2 and 3**) and because no alterations of myeloperoxidase activity in the liver of mice 10 hours after CLP was observed in response to CX43 blockade (**Supplemental Figure 6 D-E**).

Taken together these data suggest that CX43 regulates the secretion of inflammatory cytokines in an autocrine manner during sepsis by the release of ATP and activation of P2RY1.

## DISCUSSION

Sepsis remains a serious complication in modern medicine, for which specific therapies targeting the dysfunctional inflammatory reaction are still lacking (1, 18). Most of the immuno-regulatory drugs tested so far were not satisfactory in clinical settings, probably because they were blocking either the pro- or the anti-inflammatory reactions, while sepsis is a complex combination of both (19). We tested a potential means to improve sepsis outcome by targeting purinergic signaling, notably known to modulate inflammatory reactions by altering inflammatory cells activation (12).

ATP released in large quantities during inflammation in the extracellular space determines outcome after sepsis via alteration of different P2 receptor activity (8). In particular, elevated extracellular ATP drives systemic inflammation and results in tissue damage and mortality, while removal of ATP using apyrase, a synthetic ectonucleotidase, improved sepsis outcome by dampening systemic inflammatory reaction (9). However, ATP potentially plays also a protective role during sepsis, by improving killing of intracellular (via P2X7 receptor) and extracellular (via P2X4 receptor) bacteria (10, 20). Current thinking is that elevated ATP levels are the consequence of non-specific release mechanisms such as necrosis and apoptosis rather than specific controlled mechanisms (21). In the current study, we tested if specific release is critical for alteration of the inflammatory reaction and ultimately sepsis survival (11).

Our data has shown that ATP released from peritoneal macrophages is an active process that occurs in response to TLR-2 and TLR-4 agonists in a CX43-dependent manner. Increased expression of CX43 on macrophages following TLR-4 pathway activation has been previously shown (22). Our study has extended this observation by showing the functional role of CX43 for autocrine signaling in macrophages. Due to the disappearance of LPMs, SPMs are the main CX43-positive population during peritonitis, and have been shown to be highly activated to phagocytose bacteria and produce large amounts of nitric oxide (17). We have shown that both pharmacologic blockade and genetic deletion of CX43 in activated macrophages led to increased survival compared to controls. This was associated with lower levels of key inflammatory cytokines in the acute phase of sepsis including IL-6, TNF-alpha in response to CX43 blockade or deletion. Decreased levels of IL-10 and IL-33 indicate that blocking CX43 might improve the long-term outcomes, as these cytokines have been shown to be involved in immunosuppression following sepsis (23).

Decreased cytokine production by infiltrating macrophages via CX43 deletion was reversed by administration of ATPgammaS or apyrase revealing a direct impact on autocrine purinergic signaling via receptor activation or desensitization that has been shown for P2RY1 (24, 25). In turn, regulation of inflammation dependent CD39 levels were disturbed in response to CX43 deletion (26). ATP typically signals through P2 receptors in an autocrine manner. While no regulation of P2 receptors was observed, functionally relevant differences were observed. Among the P2 receptors that have been shown to play a critical role for macrophage activation (10, 27–29), we identified the P2RY1 to be critical for ATP-dependent macrophage activation. Pharmacological blockade of the CX43/ATP/P2RY1 axis at all levels led to specifically reduced cytokine secretion by isolated macrophages and ultimately to increased survival to sepsis in our MAC-CX43 KO mouse model. This effect is specific for alterations in cytokines secretion as no impairment of their phagocytic capacity was observed, adding clear evidence to previous conflicting results (30, 31).

In this study, we identified a novel key pathway for autocrine/paracrine modulation of macrophage activation that involves TLR-induced CX43 expression and ATP release. This effect is macrophage specific given the exclusive expression of CX43 on macrophages conversely to neutrophils or lymphocytes. Therefore, CX43 pharmacological blocking is likely to address macrophage function specifically.

These results are substantiated by human data where we identified CX43 positive macrophages in the peritoneal cavity of patients with peritonitis but not in control patients, allowing us to hypothesize that CX43 plays a comparable role in human macrophages than in mouse macrophages. However, as CX43 proteins form pores that are permeable for other small molecules, typically lower than 1 kDa the effect on the secretion of other nucleotides, including UDP (22), as well as other metabolites could influence sepsis outcome as well and needs to be explored in future studies.

In conclusion, we identified that CX43 is upregulated specifically on infiltrating macrophages locally and systemically during sepsis and that the elevated levels of CX43 mediates cytokine secretion but not phagocytosis of macrophages via extracellular ATP and the purinergic receptor P2Y1. Therefore, CX43 is critical for the pathophysiology of sepsis and may be a potential new target for improving the outcome of this critical clinical condition.

## METHODS

### Human samples

Peritoneal lavage fluids were collected from patients operated at the Department of Visceral Surgery and Medicine, Inselspital Bern, Switzerland (**Supplemental Table 1**). Peritoneal lavage fluid was filtered through a 100 µm filter and a Ficoll (GE Healthcare, #17-5442-02) gradient was performed by pipetting 10 ml Ficoll under 40 ml peritoneal lavage fluid in a 50 ml Falcon tube. Tubes were centrifuged (800 g, 4°C, 20 min, no brake). Cells were stained for flow cytometry as described below with anti-human antibodies (**Supplemental Table 5**).

### Experimental animals

Animals were housed in specific pathogen free (SPF) conditions. Animals used in our experiments were aged between 8 to 12 weeks and were females. WT mice were on C57Bl/6(J) background and were purchased from Harlan, Netherlands. MyD88/TRIF^-/-^ mice and caspase 1/11^-/-^ mice were provided by Andrew MacPherson (Mucosal Immunology, University of Bern) and were bred germ-free at the Clean Mouse Facility, University of Bern, Switzerland. These mice were colonized with SPF flora for experimental purposes.

Due to the fact that CX43 is expressed on various cell types including endothelial cells and myocytes in the heart in addition to inflammatory cells, CX43 full KO die perinatally due to cardiac malformation (14, 15). In order to develop a cell-specific conditional MAC-CX43 KO mouse model, B6.129P2-Lyz2tm1(cre)Ifo/J mice and B6.129S7-Gja1tm1Dlg/J mice were purchased from Jackson Laboratory (Bar Harbor, ME, USA) and were crossed together. MAC-CX43 KO mice used in our experiments were carrying a floxed Gja1 gene on both alleles and cre enzyme either on both alleles (*Lyz2^cre/cre^*, *Cx43^flox/flox^*) or on one allele for controls (*Lyz2^cre/wt^*, *Cx43^flox/flox^*). Genotyping of MAC-CX43 KO mice was done according to Jackson protocol and using following primers: Gja1 (Forward primer: 5’ CTT TGA CTC TGA TTA CAG AGC TTA A 3’ and reverse primer: 5’ GTC TCA CTG TTA CTT AAC AGC TTG A 3’) and Lyz2 (Wild-type forward primer: 5’ TTA CAG TCG GCC AGG CTG AC 3’, mutant (cre) forward primer: 5’ CCC AGA AAT GCC AGA TTA CG 3’, common reverse primer: 5’ CTT GGG CTG CCA GAA TTT CTC 3’).

### Caecal ligation and puncture

We used caecal ligation and puncture (CLP) in order to induce peritonitis leading to high-grade sepsis (32). In brief, mice were anesthetized with isoflurane and a longitudinal 1 to 2 cm midline laparotomy was performed in order to expose the caecum. The caecum was tightly ligated using a 6.0 silk suture (6-0 PROLENE, Ethicon) with 2/3 of the caecum included below the ligature and was perforated twice using an 22 gauge needle. The abdomen was closed and mice were subcutaneously injected with 1 ml of 0.9 % saline. Sham-operated mice received the same surgical operation without caecum ligation and punctures. No antibiotics were administered to mice that underwent the operation. The mice were sacrificed at 5 and 10 hours for experimental purposes. A separated set of animals, including WT and MAC-CX43 KO, were used in survival studies and were sacrificed latest 72 hours after CLP. All the animals operated with CLP were monitored using a murine sepsis score that was adapted from a previous publication (33) (**Supplemental Table 3**). Animals were treated with analgesics (buprenorphin) or sacrificed according to their score.

### Collection of blood, peritoneal fluid and organs

Prior to harvesting, mice were anaesthetized using 2 μl/kg mouse body weight of a mixed triple combination of Fentanyl (0.05 mg/ml), Dormicum (Midazolam, 5 mg/ml) and Medetomidin (1 mg/ml). Animals were placed on a surgical tray, abdominal skin was removed and laparotomy was performed. Blood was collected from inferior vena cava using a 22 gauge catheter (BD Insyte, #381223) and a 2 ml syringe into 1.5 ml Eppendorf tubes. Blood was incubated at room temperature for 60 min and centrifuged at 1’000 g for 10 min. Supernatant was transferred in a new tube and further centrifuged at 10’000 g for 10 min. Serum was finally collected in a new tube and snap frozen in liquid nitrogen. In order to collect peritoneal fluid, 5 ml ice cold PBS was injected i.p. using a 10 ml syringe and a 23 gauge needle. The abdomen was massaged for 15 seconds and at least 1 ml peritoneal fluid was collected in Eppendorf tubes. Tubes were centrifuged at 10’000 g for 10 min in order to pellet cells present in the peritoneal fluid and supernatant was pipetted into a new tube. Both pellets and supernatants were snap frozen in liquid nitrogen for further experiments. Various organs, including liver, lungs, kidney, spleen, and small intestine were collected and either further processed for flow cytometry analyses or snap frozen in liquid nitrogen for future experiments.

### Isolation and culture of primary mouse macrophages, culture of cell lines

Mice were injected i.p. with 2 ml 3% Brewer thioglycollate (Sigma-Aldrich, #B2551) 3 days prior macrophage isolation. In order to collect macrophages, the peritoneal cavity was washed with ice cold harvest medium (Dulbecco’s phosphate-buffered saline without calcium and magnesium supplemented with 3% fetal calf serum, Sigma-Aldrich, #D8537). Collected fluid was filtered through 100 µm filter and cells were centrifuged at 800 g and 4°C for 5min. Erythrocytes were lysed using erythrocyte lysis buffer (Qiagen, #160011922). Cells were counted using a counting chamber and were plated after resuspension in culture medium (Dulbecco’s Modified Eagle Medium/F12 GlutaMAX (ThermoFisher Scientific, Gibco, #10565018) supplemented with 50 IU/ml penicillin G and 50 µg/ml streptomycin and 10% fetal calf serum). HEK293-pmeLUC cells were provided by Francesco Di Virgilio (Department of Morphology, Surgery and Experimental Medicine, University of Ferrara). HEK293-pmeLUC cells were cultured in DMEM/F12 supplemented with 10% fetal calf serum, 50 IU/ml penicillin G, 50 µg/ml streptomycin and 0.2mg/ml G418 sulfate (Geneticin, Calbiochem). Cells were cultured in 75 cm^2^ tissue culture flasks (VWR, #734-2313) and kept in the incubator at 37°C and 5% CO2. The viability of cultured cells was checked under the microscope before every cell passage. After detachment of cells using a 0.05% trypsin / EDTA solution (ThermoFisher Scientific, #15400054), cells were counted using a counting chamber (Brand, #717805) and plated on 96-wells transparent flat bottomed cell culture plates (Sarstedt, Tissue culture plate 96-wells, sterile, #83.3924.300) for experimental purposes.

### Quantification of extracellular ATP

Extracellular ATP was measured using a luciferin-luciferase assay as described in manufacturer protocol (Biothema, #144-041) and bioluminescence was quantified using a Tecan Infinite 200 plate reader (Tecan). In vitro, the supernatant was collected from 96-wells cell culture plate. The ecto-ATPase inhibitor ARL 67156 (Sigma-Aldrich, #67156) was used in order to reduce extracellular ATP degradation. In vivo, extracellular ATP was quantified in the peritoneal fluid and in the plasma using the same kit and 1:20 dilution in PBS. Plasma was collected in Eppendorf tubes with CTAD (citrate-theophylline-adenosine-dipyridamole) (BD Vacutainer, #367599) in order to avoid platelet activation and ATP release via degranulation.

### In vivo ATP imaging using plasma-membrane targeted luciferase

Measurements of extracellular ATP using a chimera plasma-membrane targeted luciferase on HEK293 cells were performed as previously described and by adapting the protocol for our own purposes (34, 35). Cells were cultured as described above. 5×10^6^ cells were injected i.p. right after CLP or sham and before suturing the abdominal wall. 6 hours after CLP, 3 mg D-luciferin diluted in 100 µl PBS were injected i.p. Bioluminescence was measured using a gamma camera (NightOWL LB 983, Berthold Technologies) 8 minutes after D-luciferin injection and is proportional to the amount of ATP present in the peritoneal cavity.

### Quantification of Colony forming units (CFUs) in blood and peritoneal lavage

As previously described (10), blood and peritoneal lavage fluid were diluted serially in sterile physiologic saline. Fifty microliters of each dilution was aseptically plated and cultured on LB agar or trypticase blood agar plates (BD Biosciences) at 37°C. After 12–16 h of incubation, the number of bacterial colonies was counted. The number of cultures is expressed as CFUs per milliliter of blood or peritoneal lavage fluid.

### Gene expression analysis

Total RNA was isolated from tissue by TRIzol reagent and following manufacturer’s protocol (Roche, #11667165001). RNA concentration and quality was analysed by spectrophotometer NanoDrop ND-1000 (Thermo Scientific). 500 ng of total RNA was used for cDNA synthesis. cDNA was synthesized by using Omniscript RT Kit 200 (Quiagen, #205113). qPCR analysis was performed by using TaqMan gene expression assays (Applied Biosystem) according to manufacturer’s protocol. mRNA was analysed by reverse transcriptase-quantitative PCR (ABI 7900, SDS 2.3 software). Primers and probes sequences were purchased from Applied Biosystems: *Arg1* (Mm00475988_m1), *Cd39/Entpd1* (Mm00515447_m1), *Cd73* (), *Il-10* (Mm00439614_m1), *Inos* (Mm00440485_m1), *Il-12rb2* (Mm00434200_m1), *P2rx1* (forward: 5’ ACT GGG AGT GTG ACC TGG AC 3’, reverse: 5’ TCC CAA ACA CCT TGA AGA GG 3’), *P2rx2* (forward: 5’ CAA AGC CTA TGG GAT TCG 3’, reverse: 5’ CCT ATG AGG AGT TCT GTT 3’), *P2rx3* (forward: 5’ ATT AAG ATC GGC TGG GTG TG 3’, reverse: 5’ TCC CGT ATA CCA GCA CAT CA 3’), *P2rx4* (forward: 5’ TAC GTC ATT GGG TGG GTG TT 3’, reverse: 5’ CTT GAT CTG GAT ACC CAT GA 3’), *P2rx5* (forward: 5’ GGG GTT CGT GTT GTC TCT GT 3’, reverse: 5’ CAC TCT GCA GGG AAG TGT CA 3’), *P2rx6* (forward: 5’ GTG GTA GTC TAC GTG ATA GG 3’, reverse: 5’ GCC TCT CTA TCC ACA TAC AG 3’), *P2rx7var1-3* (forward: 5’ CTT GCC AAC TAT GAA CGG 3’, reverse: 5’ CTT GGC CTT TGC CAA CTT 3’), *P2rx7var4* (forward: 5’ TCA CTG GAG GAA CTG GAA GT 3’, reverse: 5’ TTG CAT GGA TTG GGG AGC TT 3’), *P2ry1* (forward: 5’ TTA TGT CAG CGT GCT GGT GT 3’, reverse: 5’ CGT GTC TCC ATT CTG CTT GA 3’), *P2ry2* (forward: 5’ GAG GAC TTC AAG TAC GTG CT 3’, reverse: 5’ ACG GAG CTG TAA GCC ACA AA 3’), *P2ry4* (forward: 5’ AAC AAC TGC TTC CTC CCT 3’, reverse: 5’ AAG TCC TAG AGG TAG GTG 3’), *P2ry6* (forward: 5’ CCT GAT GTA TGC CTG TTC AC 3’, reverse: 5’ CAC AGC CAA GTA GGC TGT CT 3’), *P2ry11* (forward: 5’ TGT GGC CCA TAC TGG TGG TTG AG 3’, reverse: 5’ GAA GAA GGG GTG CAC GAT GCC CA 3’), *P2ry12* (forward: 5’ ATA TGC CTG GTG TCA ACA CC 3’, reverse: 5’ GGA ATC CGT GCA AAG TGG AA 3’), *P2ry13* (forward: 5’TGC AGG GCT TCA ACA AGT CT 3’, reverse: 5’ CCT TTC CCC ATC TCA CAC AT 3’), *P2ry14* (forward: 5’ GGA ACA CCC TGA TCA CAA AG 3’, reverse: 5’ TGA CCT TCC GTC TGA CTC TT 3’). Relative changes in mRNA were calculated with the ΔΔCt method. Ct values of target genes was calculated relative to a reference control gene (*Tbp* (Mm01277042_m1) and Beta-actin (*Actb*, Mm00607939_s1)) using the following formula ΔCtTG= CtTG-CtRG. Experimental groups are normalized to control group. ΔΔCt=ΔCtexp - ΔCtcon. Fold change=2-ΔΔCt Log2FC.

### Immunoblot

Proteins were extracted using RIPA Lysis buffer, which volume was adapted for every tissue. Protein extracted were quantified using Bio-Rad Protein Assay System (Bio-Rad Laboratory, Melville, NY). Protein lysates were boiled in Laemmli buffer, run on BIO-RAD Mini-Protean^®^ TGXTM (10–20% Ready Gel^®^ Tris-HCl Gel System, 12-well comb #456-1095 and 10-well comb #456-1094) for 90 minutes at 120 V, and transferred on BIO-RAD Trans-Blot^®^ Turbo™ Mini or Midi PVDF membranes (Mini PVDF Transfer Packs #170-4156 and Midi PVDF Transfer Packs #170-4159) by semi dry Transfer (Trans-Blot^®^ Turbo™ Transfer System BIO-RAD). Specific proteins were detected using the following primary antibodies: Connexin-43 (1:850) (Cell Signalling, #3512), β-actin HRP-conjugated (1:5000) (Sigma-Aldrich, #2228). Antibodies were diluted in 5% milk and incubation was done overnight at 4°C. Following secondary antibodies were used: HRP-conjugated anti-rabbit (1:5000) (Dako, #P0448). Membranes were then washed and protein expression was analysed by chemiluminescence (Western Lightning Plus-ECL Perkin Elmer), using Fusion-FX (Vilber).

### Immunohistochemistry and Immunofluorescence

CX43 expression in the liver and in the lungs was assessed by immunostaining. In brief, paraffin embedded tissue sections were dewaxed followed by incubation in antigen retrieval citrate buffer (Sigma-Aldrich, #C9999). Erythrocyte quenching was performed using 3% H_2_O_2_ (Dr. Grogg, #K45052100407) diluted in phosphate buffered saline supplemented with 0.05% Tween 20 (PBST). After protein blocking (PBS, 5% BSA), the slides were incubated overnight at 4°C with the primary rabbit anti-CX43 antibody (1:100, Cell Signalling, #3512), followed by incubation with secondary biotinylated anti-rabbit (Dako, #E0432) for 1h at room temperature and by incubation with peroxidase labelled streptavidin (Seracare, #71-00-38) for 30 min at room temperature. Liver and lungs sections were stained with hematoxylin (MERK, #HX43078349) for 1 min followed by Eosin staining for 6 min. Later sections were mounted with Eukritt (Kindler, GmbH).

Primary peritoneal macrophages were isolated as described above and plated on a chamber slide for immunofluorescence. After incubation at 37°C for 12 hours, cells were washed using PBS and fixed using 4% formaldehyde solution for 15 minutes. Specimens were covered with blocking buffer (1X PBS / 5% normal serum / 0.3% Triton™ X-100) for 60 minutes followed by overnight incubation with primary rabbit anti-CX43 antibody (1:100, Cell Signalling, #3512). Further, specimens were incubated 120 minutes with secondary goat anti-rabbit IgG FITC-conjugated antibody (Sigma, #F0382) and 5 minutes in Dapi (Sigma, #D9542). Finally, specimens were mounted using Vectashield Antifade Mounting Medium (Vector Laboratories, #H-1000) and covered with a coverslip.

### Cytokine measurement

Inflammatory cytokines (TNF-alpha, IL-6 and IL-10) were quantified in serum, peritoneal fluid and organs collected in mice following CLP and in control mice, and supernatants from in vitro LPS-stimulated peritoneal macrophages. Uncoated ELISA kits were purchased from Invitrogen (Thermo Fisher Scientific) and ELISA were performed following manufacturer’s protocol: TNF-alpha (Invitrogen, #88-7324-22), IL-6 (Invitrogen, #88-7064-22), IL-10 (Invitrogen, #88-7105-22). Plates were red using Tecan.

Serum cytokines were measured using electrochemiluminescence assays (Meso Scale Discovery (MSD), Rockville, Maryland, USA). Serum TNF-alpha, IL-6, IL-10 and IL-33 were measured using a multiplex assay using the V-PLEX Mouse Cytokine 19-Plex kit (MSD). Samples were diluted 1:3 in proprietary buffer (MSD) and measured in duplicate according to manufactures protocol. Plates were run on MSD plate reader model 1250.

### Neutrophil detection by myeloperoxidase assay

Myeloperoxidase assay was carried out as previously described (36, 37). Briefly, liver samples were thawed, 50 mg of tissue was weighed out and homogenized in 1 ml of 20 mM phosphate buffer using a TissueLyser (Qiagen) 2 min at 20 Hz, centrifuged at 10,000 g for 10 min, and the pellet was resuspended in 50 mM phosphate buffer (pH 6.0) containing 0.5% hexadecyltrimethyl ammonium bromide. The samples were then freeze-thawed four times using liquid nitrogen followed by a 40 sec sonication, centrifuged at 10,000 g for 10 min, and the supernatant was used for the MPO assay. The assay mixture contained 10 µL of the sample, 25 µL 3,3’5,5-tetramethylbenzidine (final concentration 1.6 mM), 25 µL H_2_O_2_ in 80 mM phosphate buffer (pH 5.4, final concentration H_2_O_2_ 0.3 mM), and 40 µL of 50 mM phosphate buffer (pH 6.0) containing 0.5% hexadecyltrimethyl ammonium bromide. The mixture was incubated at 37°C for 2 min, stopped with the addition of 1 M HCl, and the absorbance was measured at 450 nm.

### Flow cytometry

To perform flow cytometry, leucocytes were isolated from the peritoneal cavity, the liver. Isolation of peritoneal leucocytes was performed as described above by washing the peritoneal cavity with ice cold PBS. The liver and the lungs were cut in small pieces and placed in 10 ml RPMI containing 3% FCS 1 mg/ml collagenase IV and 0.1 mg/ml DNase I (Roche) with shaking at 37°C for 30 min. Cell suspension was passed through a 100 µm cell strainer and washed twice with FACS buffer (PBS, 3% FCS, 2 mM EDTA). Cells were centrifuged (800 g, 7 min, 4°C) and resuspended in 40% Percoll solution and layered on top of 80% Percoll solution. Gradient centrifugation was carried out (900 g, 20 min, 4°C, no brake). Leucocytes were collected from the interphase, washed with FACS buffer and centrifuged (800 g, 7 min, 4°C). Cells were finally resuspended in FACS buffer and further stained for flow cytometry.

Aliquots of 10^6^cells/100 μL of staining buffer per well were incubated each with 1 μg of purified anti-CD16/CD32 for 20 min at 4°C in the dark, in order to block non-specific binding of antibodies to the FcγIII and FcγII receptors. Cells suspension were incubated with a fixable viability dye (AmCyan, eBioscience, #65-0866-14) diluted in DPBS during 20 min at 4°C in the dark to exclude dead cells. Subsequently, these cells were separately stained with the surface markers for 20 min at 4°C in the dark with 1 μg of primary antibodies. For cytokines and transcription factors, cells were stained with antibodies to surface antigens, fixed and permeabilized according to the manufacturer’s instructions (Foxp3/Transcription Factor Staining Buffer Set; eBioscience, 00-5523-00). Fluorescently labeled anti-mouse used are summarized below (**Supplemental Table 4**). Cells were subsequently washed twice with, and resuspended in FACS buffer. Finally, cell data were acquired on a LSR II SORP H271 (BD Biosciences). Flow cytometric analysis was done using FlowJo (Treestar). In all experiments, FSC-H versus FSC-A was used to gate on singlets with dead cells excluded using the fluorescence-coupled fixable viability dye. Murine inflammatory peritoneal cells were gated as follow: (1) SPMs: viable (AmCyan^low^), CD45^high^, CD3 and CD19^low^, Ly6G^low^, F4/80^int^ and CD11b^int^ (2) LPMs: viable (AmCyan^low^), CD45^high^, CD3 and CD19^low^, Ly6G^low^, F4/80^high^ and CD11b^high^ (3) Neutrophils: viable (AmCyan^low^), CD45^high^, CD3 and CD19^low^, Ly6G^high^, F4/80^low^ and CD11b^low^ (4) Dendritic cells: viable (AmCyan^low^), CD45^high^, CD3 and CD19^low^, Ly6G^high^, F4/80^low^, CD11b^low^ and CD11c^high^. Liver macrophages were characterized as published previously (38). Inflammatory cells in the liver were gated as follow: AmCyan^low^, CD45^high^, CD3/CD19^low^.

Human peritoneal macrophages were gated as follow: AmCyan^low^, CD45^high^, CD3^low^, CD19^low^, CD56^low^, CD11c^low^, CD16^high^ and CD67^low^.

### Time-of-flight Mass Cytometry (CyTOF)

For time-of-flight mass cytometry (CyTOF) experiments, cells were isolated from the peritoneal cavity as previously describe (39), 3×10^6^ cells were stained with lanthanide-conjugated antibodies as previously described (40). Antibodies were purchased already labelled or labelled using the MaxPar antibody labelling kit (Fluidigm, **Supplemental Table 6**). Raw signals were normalized to median bead intensity using four elemental calibration beads (Fluidigm) as previously described (41). The Live intact single events were gated for their DNA content (MaxPar Intercalator, Ir 193) and viability (Cell-ID Cisplatin, 198 Pt). CD45^neg^ cells were excluded. Analysis was done using t-SNE for dimensionality reduction on arc sinus transformed data (42). On t-SNE plots the clusters were manually gated based on their marker expression.

### Phagocytic assay

The phagocytic capacity of plated macrophages was assessed as described in the manufacturer protocol (Cayman Chemical, #500290). Peritoneal macrophages were isolated from WT and MAC-CX43 KO mice as described above. 50’000 macrophages were plated per well on a chamber slide (Nunc Lab Tek II, #154534) and incubated for 12 hours at 37°C and 5% CO_2_. Macrophages were primed or not with LPS 1µg/ml for 4 hours and exposed or not to FITC latex beads coated with rabbit IgG diluted 1:200 for 2 hours (4 groups in total). Negative controls were incubated on a separate slide chamber at 4°C.

### Resazurin cell viability assay

Cell viability after stimulation with different agonists was assessed using resazurin assay (Sigma-Aldrich, #199303). Cells were exposed to a 1.4 μM resazurin solution and fluorescence was quantified using Tecan-Reader^®^.

### Compounds

Toll-like receptors 4 and 2 agonists were used to stimulate peritoneal macrophages in vitro: lipopolysaccharides (LPS) from E.coli O111:B4 (Sigma-Aldrich, # L4130) and Pam3CSK4 (Invivogen, # tlrlpms). In order to block ATP release via hemichannels, 18-alpha-glycyrrhetinic acid (Sigma-Aldrich, # G8503) was used as a nonspecific inhibitor of connexin channels and Gap27 (Tocris Bioscience, # 1476) was used as a specific CX43 inhibitor. Probenecid (Sigma-Aldrich, # P8761) was used as a specific pannexin-1 inhibitor and carbenoxolone (Sigma-Aldrich, # C4790) as a non-specific inhibitor of connexin and pannexin channels.

### Statistical analysis

Survival statistics were determined by the Kaplan-Meier curve and log-rank test. t-test and ANOVA were used to assess the significance of the differences in the means of the different populations. When necessary, *p*-values were corrected for multiple comparisons using Turkey correction. Statistical tests were performed using Prism 6 software (GraphPad Software). Levels of significance were assessed with specifically indicated tests and *p*-values are presented as follows: ns *p* non-significant; * *p* < 0.05; ** *p* < 0.01; *** *p* < 0.001 and **** *p* < 0.0001. t-SNE visualizations were produced with R-package Rtsne on arc sinus transformed data (43).

### Study approval

All human studies were approved by the Ethical Commission of the Canton Bern and written informed consent was obtained from all subjects. Peritoneal fluid collection at the beginning of an operation was included in a larger clinical trial, whose protocol is published on ClinicalTrials.gov (NCT03554148, Study ID Number: Bandit).

Animal experiments were planned, carried out and reported in agreement with current 3R and ARRIVE guidelines (44) and approved according to Swiss animal protection laws by the Veterinary Authorities of the Canton Bern, Switzerland (license no. BE 4/15).

## AUTHOR CONTRIBUTIONS

M.D. and G.B. designed research studies.

M.D. performed the experiments described in the publication.

N.M. provided help in conducting experiments, acquiring data and analyzing data.

N.M. and F.J. provided help for flow cytometry experiments.

J.Z. provided help for mass cytometry experiment.

D.Sa. provided help for tSNE analysis and statistical analyses.

M.D. and G.B. wrote the manuscript.

D.St., D.C. and G.B. supervised the studies and reviewed the manuscript.

## ACKNOWLEDGMENTS

The authors would like to thank collaborators from the Laboratory of Visceral and Transplantation Surgery, Department for BioMedical Research, University of Bern for technical assistance. Mass cytometry was performed at the Cytometry Facility of the University of Zurich.

